# Gene expression as phenotype - Many small-step changes leading to little long-term phenotypic evolution

**DOI:** 10.1101/2022.06.24.497468

**Authors:** Pei Lin, Guang-An Lu, Zhongqi Liufu, Yi-Xin Zhao, Yongsen Ruan, Chung-I Wu, Haijun Wen

## Abstract

Unlike in genotypic evolution, there are few general rules governing phenotypic evolution with one of them being the small-step evolution. More specifically, natural selection tends to favor mutations of smaller phenotypic effects than of larger ones. This postulate can be viewed as a logical extension of Fisher’s Geometric Model (FGM). Testing this FGM postulate, however, is challenging as the test would require a large number of phenotypes, each with a clear genetic basis. For such a test, we treat the expression level of each gene as a phenotype. Furthermore, a mechanism of small-step expression evolution exists, namely via the control by microRNAs (miRNAs). Each miRNA in metazoans is known to weakly repress the expression of tens or hundreds of target genes. In our analysis of mammalian and Drosophila expression data, small step evolution via miRNA regulation happens frequently in long-term evolution. However, such small-step evolution does not lead to long-term phenotypic changes which would take too many such steps to accomplish. Furthermore, target site changes often cancel themselves out by continual gains and losses. The results suggest that the FGM postulate may be most appropriate for phenotypic fine-tuning near the expression optimum. In contrast, longterm expression evolution may occasionally take large steps (e.g., mutations in transcription factors) when big environmental shift happens. In another study (Lu et al. 2021), we further show how the small-step evolution of expression phenotypes is a manifestation of miRNAs’ role in developmental canalization. In conclusion, the rules of phenotypic evolution may depend crucially on the genetics of the phenotype, rather than its metric properties.

## Introduction

With the massive applications of DNA and RNA sequencing, the understanding of genotypic evolution has made large leaps in recent years. The next frontier of phenotypic evolution may be a much more challenging task, due to the lack of common rules such as those widely invoked in genotypic evolution (Wright 1931; Kimura 1968; Li et al. 1981). Nevertheless, there is a common postulate about phenotypic evolution in that it proceeds in small steps (Fisher 1930). Indeed, a premise of Darwinian evolution seems to be gradualism while large-step evolution is often attributed to cumulative small-step changes over a period of time (Futuyma 2013). Skeletal evolution in sticklebacks is a recent example of such studies (Miller et al. 2014).

In discussing phenotypic evolution, it would be necessary to define the step size of phenotypic change. Here, we focus on the phenotypic effects of individual mutations. The step-size of would then be determined by the class of mutations of interest. For example, we may ask whether a mutation in transcription factors would lead to a larger phenotypic change (say, the wing size of insects) than in structural proteins. An explicit example is the substitution between two amino acids and the step size would be the physico-chemical distance between them (Chen et al, 2019a, 2019b; Bergman and Eyre-Walker 2019). Interestingly, the conclusion of Chen et al 2019b is that the selective advantage tends to be associated with big-step changes. However, since the physico-chemical distances between amino acids are reflected by their evolutionary potentials (which are weakly correlated to many actual physico-chemical distances; see Chen et al. 2019a), we now search for phenotypes that can be directly and abundantly measured such that we can analyze the phenotypic evolution in an appropriate theoretical framework.

### Fisher’s Geometric Model and its postulate

The best-known mutation-based model of phenotypic evolution may be Fisher’s Geometric Model (FGM, Fisher 1930; Tenaillon 2014). In FGM, a mutation that has a smaller phenotypic effect is more likely to be beneficial than one with a larger effect. It is easier to picture FGM in a three-dimensional space with multiple fitness peaks on a 2D terrain (although FGM is based on a multi-dimensional fitness landscape (Wagner and Zhang 2011)). When a population is near, but not right on top of, the fitness peak, any large displacement is likely to overshoot the peak, resulting in fitness reduction. In contrast, as the displacement becomes very small, the chance of going up toward the peak would approach 0.5.

We now refer to the view that “smaller-step phenotypic changes are more likely to be beneficial than large-step changes” as the FGM postulate. Although studies have applied FGM to various evolutionary questions with or without invoking this postulate, a rigorous test of the FGM postulate would indicate whether FGM itself is suited as a phenotypic model. While FGM has been widely used to study adaptation, quantitative genetics and deleterious mutations (Orr et al. 1998; Sellis et al. 2011; Lourenço et al. 2013; Matuszewski et al. 2014; Huber et al. 2017; Simons et al. 2018), this study may be the first one aimed at testing the FGM postulate directly. For a more comprehensive review and commentaries on previous studies of FGM, please refer to the Supplementary note.

### Gene expression, microRNAs and the FGM postulate

In the attempt to test the FGM postulate, we note that few phenotypic changes have been sufficiently well-mapped to reveal even a single underlying mutation, let alone the many mutations that may collectively drive the long-term phenotypic evolution. In this context, gene expression would be a good phenotype for testing the FGM postulate. First, the expressions of thousands of genes between species have been measured. Second, a class of regulators of gene expression is known as microRNAs (miRNAs) that down-regulate the expression of their target genes (Bartel 2004; Bartel 2009; Lu et al. 2008; Wu et al. 2009; Tang et al. 2010; Shen et al. 2011; Hausser and Zavolan 2014; Lyu et al. 2014). In any tissue, the most highly expressed 50 miRNAs may account for > 95% of the miRNA abundance (Wen et al. 2014 and Supplementary Table 1). Third, each miRNA regulates the expression of hundreds of target genes by binding to the target sites on their 3’ UTR. It is estimated that more than 30% of expressed genes are regulated by miRNAs (Xu et al. 2013; Agarwal, et al. 2015). Hence, the expression evolution can be tracked by examining the gains and losses of these target sites between species.

In contrast with mutations in transcription factors that may alter gene expression by several hundred percent (Royzman et al. 1997; Guarner et al. 2017; Lambert et al. 2018), the miRNA effect on gene expression is generally modest, typically changing expression by no more than 10% (Chen et al. 2019). Such phenotypic effects are often smaller than the variation of gene expression in natural populations (Eichhorn et al. 2014; Liufu et al. 2017; Zhao et al. 2017; Lu et al. 2018a, 2018b; Chen et al. 2019; Zhao et al. 2021). In this sense, the evolution in the miRNA target sites may be an ideal system of small-step phenotypic evolution. We then wish to know whether these small-step changes would contribute cumulatively to long term expression evolution.

In brief, we propose that the rules of phenotypic evolution may depend on the underlying genetics of the phenotypes of interest. Gene expressions may be such phenotypes with respect to miRNA regulation. Lastly, in another study we knocks down the Dicer gene that is critical in miRNA biosynthesis (Lu et al. 2021). By doing so, we reduce the expressions of hundreds of miRNAs, but only mildly. The resulting phenotypic changes thus shed considerable light on the functional basis of the FGM postulate.

## Results

To answer the question of whether small-step changes take place frequently in phenotypic evolution, we examine the evolution of miRNA target sites in relation to gene expression. Nevertheless, the first step would have to be about the conservation of the regulators, i.e., the miRNAs, themselves.

### 1. Conservation of the expression of miRNAs

In studying the evolution of miRNA target sites, it would be desirable to use miRNAs of comparable abundance. By doing so, we ensure that the expression changes in the target genes are due to target site evolution, rather than miRNAs themselves. (Here, each miRNA is in fact a family of miRNAs that share the same seed and regulate the same set of targets.) For each tissue in each species, we use the most highly expressed 20 miRNAs that generally account for >= 85% of the sequencing reads (Supplementary Table 1). The list of top-20 miRNAs are largely overlapped between species (Fig. 1). For example, the heart tissue of human and mouse share 16 of the 20 miRNAs (hence, there are 24 = 16 + 4*2 data points). On average, we consider 26 miRNA families in each tissue (with an average of 2060 target genes; Table 1). These miRNAs, being highly correlated in their expression between species, allow us to focus on the divergence of their target genes.

**Figure. 1.**
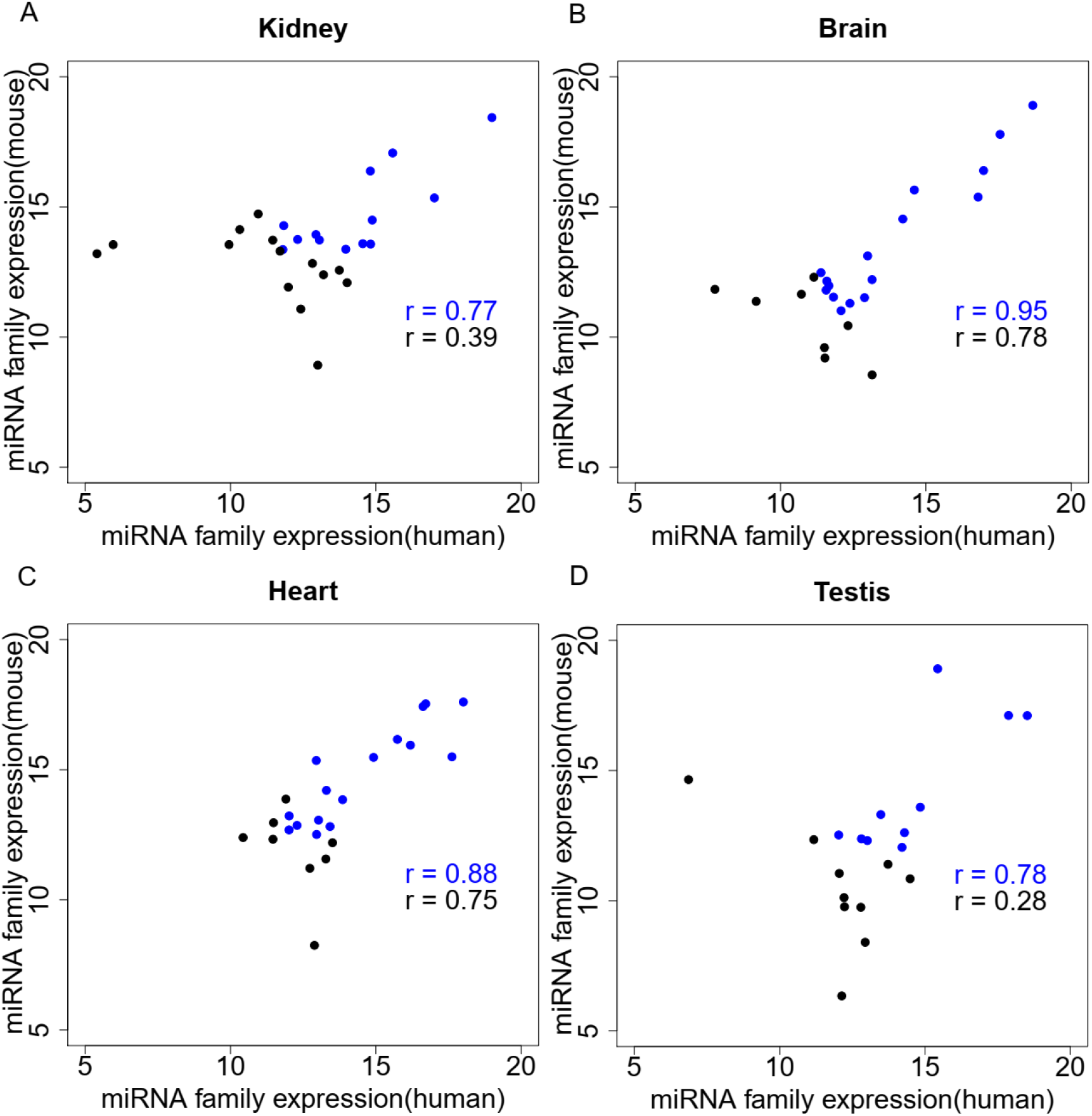
The high correlation of miRNA expressions between human (X-axis) and mouse (Y-axis) ensures that the target gene expression differences between human and mouse are due to target site, rather than miRNA, changes. Each dot represents a miRNA (or a family of miRNAs with identical seeds) with blue dots representing those in the top-20 in both human and mouse. Black dots represent those that are among top-20 in one species only. Pearson correlation coefficients (r) are shown. Number of reads per million (log2 transformed) is used for measuring miRNA family expression.

**Table 1.** miRNA target site effect on gene expression evolution between human and mouse.

| Tissue | Number of genes | Observed |  |  | Knock-out based prediction |  |
| --- | --- | --- | --- | --- | --- | --- |
| | | $R^2$ | Slope | p-value | $R^2$ | Slope |
| **Kidney** | 2743 | 0.03 | -0.08 | $< 1e-15$ | 0.099 (0.068,0.137) | -0.091 (-0.10,-0.07) |
| **Brain** | 2284 | 0.01 | -0.03 | $< 1e-05$ | 0.106 (0.067,0.149) | -0.091 (-0.10,-0.07) |
| **Heart** | 1876 | 0.02 | -0.06 | $< 1e-09$ | 0.096 (0.054,0.14) | -0.091 (-0.11,-0.07) |
| **Testis** | 1337 | 0.01 | -0.06 | $< 1e-03$ | 0.09 (0.045,0.144) | -0.091 (-0.11,-0.06) |

NOTE. Repression strength per target site observed in microRNA KO experiments is used to predict gene expression change. 95% confidence interval are listed inside parentheses.

### 2. Assessing the repression strength of miRNAs in the deletion experiments

To assess the strength of target site changes on gene expression, we collected expression data from miRNA deletion lines of mice and Drosophila. The expression changes due to miRNA knockout will be used in a later section on the expected expression differences between human and mouse due to target site evolution. The three chosen mouse miRNAs (miR-1, miR-122, and miR-181) are all highly expressed (among the top-20) and strongly conserved across vertebrates. The three Drosophila miRNAs are also conservative and expressed at moderate to high level. As expected, target genes are significantly de-repressed upon miRNA knock-out (Figure 2). Conserved target sites generally exert a stronger effect than non-conserved sites (Supplementary Table 2). Note that the knock-out experiment of liver-specific miR-122 was performed in the mouse and the expression was assayed in the mouse liver.

**Figure 2.**
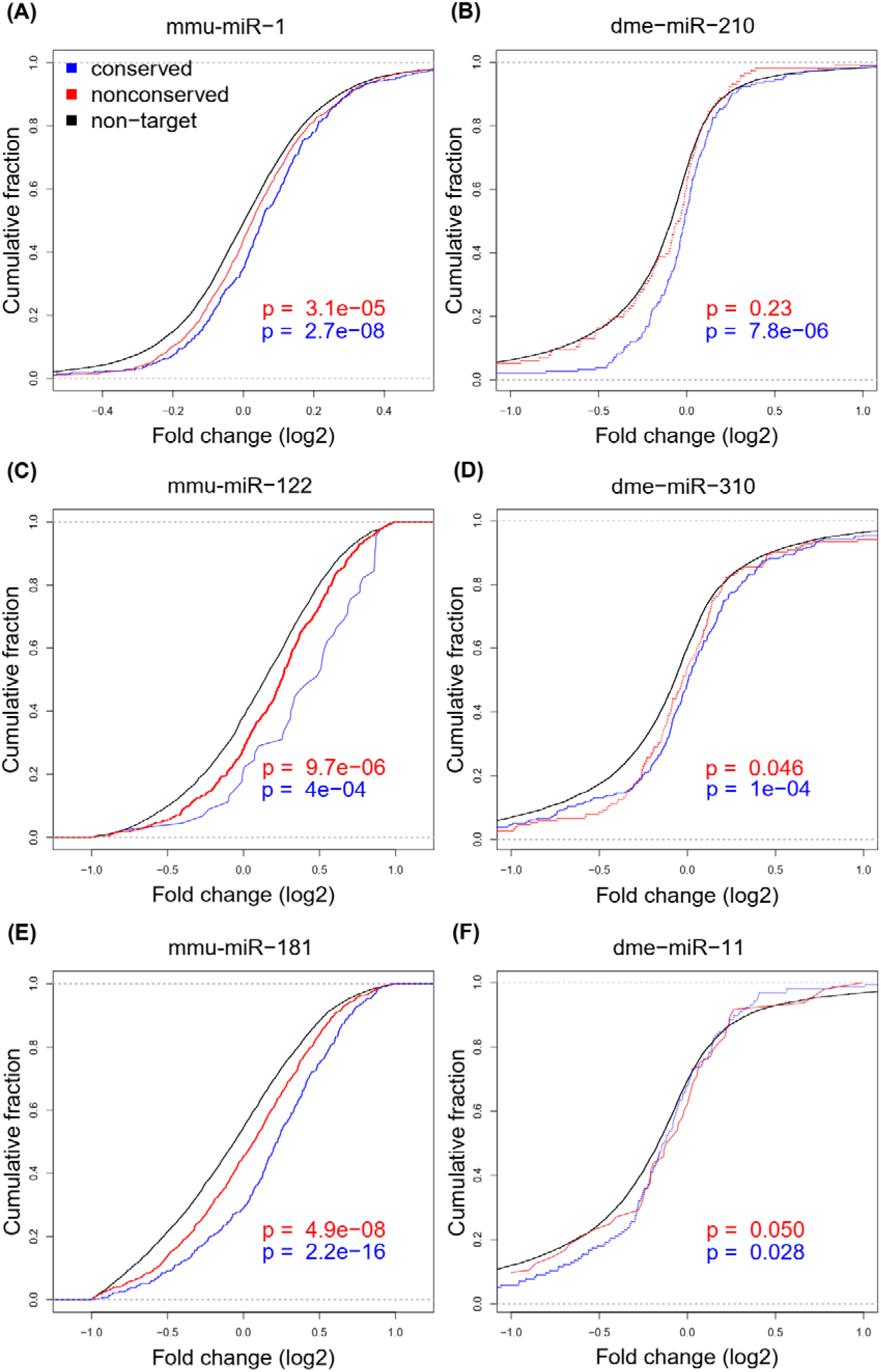
microRNAs exert weak repression on target gene expression. Cumulative distributions are shown for six conservative miRNAs (mmu for mouse-human and dme for D. melanogaster - D, yakuba comparison).

A previous study (Chen et al. 2019) reported the median repression by miRNAs to be only 8% when many miRNAs are considered. The repression strength of an average miRNA is said to be not much larger than the natural variation in the expression level (Lu et al., [the companion study and the references cited therein]). Our analyses corroborate the consensus that the repression strength of miRNAs is weak to moderate. In this respect, the knockout effect of miR-122 is unusually strong, likely due to the high abundance of miR-122, as well as the small number of conserved target genes expressed in the liver (< 100 genes).

Target sites were identified using TargetScan (7mer-A1, 7mer-m8, and 8mer-A1, see methods). Blue curves represent genes with at least one conservative target site and red curves are for genes with the target site in mouse only or in D.melanogaster only. Fold change of expression level for each gene was determined by comparing KO to control. For each miRNA KO experiment, target genes are significantly up-regulated when compared with background genes (the Kolmogorov–Smirnov test).

### 3. Changes in miRNA target sites are frequent in evolution

We now evaluate the evolution in gene expression by analyzing target site changes. Let Hi be the number of target sites regulated by the top-20 miRNAs in gene i in human but not in mouse. Target sites present in primates (human, chimpanzee and macaque) and absent in rodents (mouse and rat) are counted. Similarly, Mi is the corresponding number in mouse and rat, but absent in primates. Note that the number of target sites in each species would be Hi + Bi or Mi + Bi (Bi being the number of sites in both species). For Drosophila species, we compare the D. melanogaster - D. simulans - D. schellia clade against the D. yakuba - D. erecta clade. Two such examples of species comparison are given in Fig. 3A and 3B.

**Figure 3.**
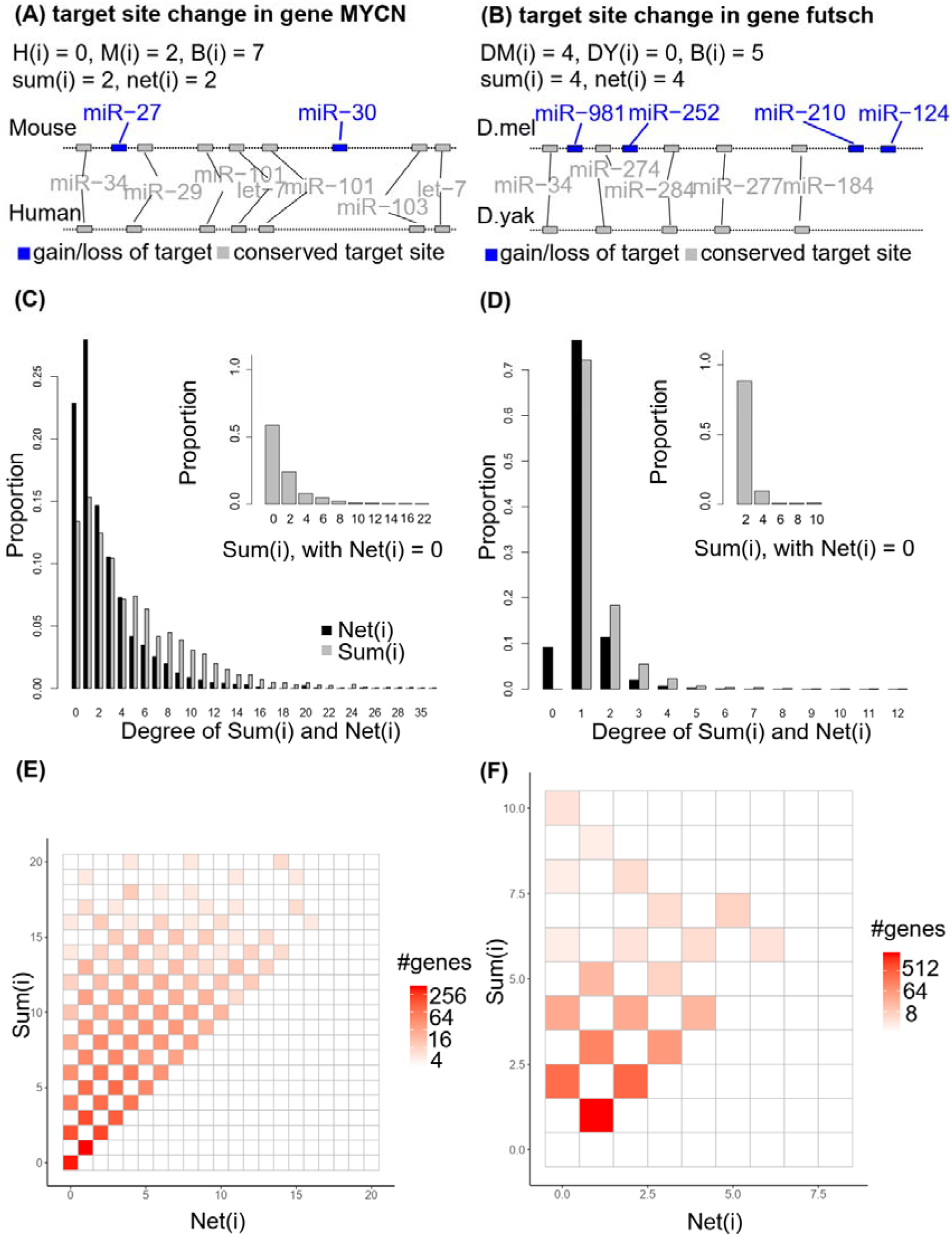
frequent gains and losses of miRNA target sites during evolution. **(A and B)**: two examples for the calculation of sum(i) and net(i). Blue and grey rectangles represent divergent and conserved sites, respectively. **(C and D)**: distribution of sum(i) and net(i); Inset in each panel is the distribution of sum(i) when net(i) = 0. **(E and F)**: number of genes with specific degree of sum(i) and net(i). Human - mouse comparisons are shown in A,C and E and Drosophila comparisons are shown in B, D and F.

The actual evolutionary changes should be at least

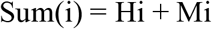

While the net number of observable changes in evolution is

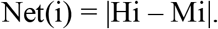

Obviously, Sum (i) should be no smaller than Net(i). Figure 3C shows the distribution of Sum(i), which is often > 5 whereas Net(i) is often < 3. The inset of Fig. 3C shows that more than 40% of genes with Net(i) = 0 have Sum(i) > 2. These genes have no extant difference in their cumulative miRNA mediated repression, given Net(i) = 0. Nevertheless, small changes in expression may not be uncommon between mouse and human as Sum(i) often exceeds 2. A similar conclusion is reached for Drosophila (Figure 3D). Net(i) and Sum(i) in Drosophila are both smaller than those in human, likely due to the much shorter 3’UTR in Drosophila.

Figures 3E and 3F show the relationships between Sum(i) and Net(i). On the diagonal are the number of genes with Sum(i) = Net(i). The total number of genes above the diagonal accounts for > 50% of all genes for both mammals and Drosophila. In many genes, Sum(i) is larger than Net(i) by more than 10. Note that Sum(i) may still under-estimate the length of the evolutionary path as gains/losses at the same site would not have been detected. Furthermore, the number of active miRNAs in a tissue can be more than a hundred, but this study is restricted to the top-20 highly expressed miRNA families. These target genes of lowly expressed miRNAs may experience numerous small-step changes.

If small-step changes were deployed to realize long term expression evolution, continual gains (or continual losses) of microRNA target sites should occur in a given lineage. In other words, we may observe target site changes tend to be all-present, or all-absent, in a given species and Sum(i) should be close to Net(i). The observation of Sum(i) > Net(i), nevertheless, is common. A simple explanation is that each gene frequently loses and gains target sites in response to small environmental changes without persistent changes in either direction. The excess of Sum(i) over Net(i) could be due to the fine-tuning of gene expression near an optimum. The analysis suggests that the evolutionary path between species in terms of small-step expression changes is quite long (Supplementary Table 3). However, they do not account for much of the net amount of evolution in the long run.

### 4. The long term expression evolution in relation to Net(i)

The key element of Fig. 3 is the Sum(i) values, i.e., the observable length of evolution between human and mouse. While Net(i) does not reflect the actual amount of evolution, it does reflect the current states of the species. As species fine-tune their optimal expression level, they may happen to differ by Net(i) in the i-th gene at present. We may thus ask if Net(i) impacts the differences in gene expression between the extant human and mouse.

More generally, the question is “between distantly related species, do the *mean* expression levels differ by an amount expected by miRNA target site changes?” Indeed, based on the miRNA knockout experiments (see Fig. 2), the expected changes can be formulated. We then ask the second question: how much of the *variation* in the expression level among genes is explainable by these target site changes? Since many factors, such as transcription factors, could influence the expression level more strongly than miRNAs can, the explainable proportion may be small. To answer these two questions, Fig. 4 plots the observed expression divergence against Net(i) between human and mouse. The slope and the correlation in these plots would answer, respectively, the first and the second question.

**Figure 4.**
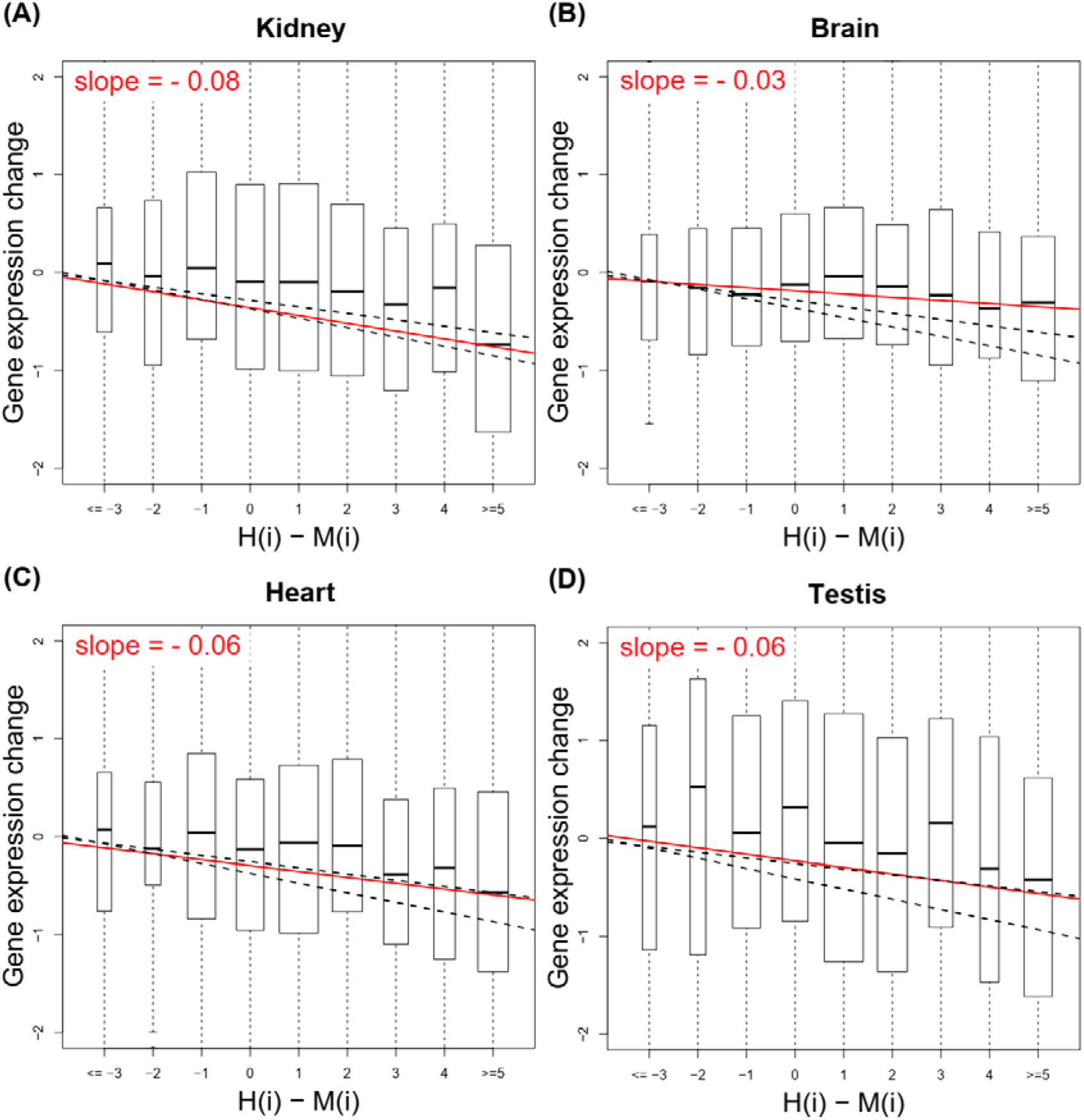
Weak contribution to gene expression divergence by the net gains of microRNA target sites. Observed target gene expression divergence was regressed on net change of miRNA target sites, which was measured as H(i) – M(i). Note that log2 transformed expression change was shown, which comparing human to mouse. Resultant slopes (red) indicate the effect size of miRNA target site while black dashed lines indicate the 95% confidence interval, derived by simulating target gene expression change driven by miRNA regulation alone.

The slope is indeed negative between gene expression level and Net(i). The regression analysis shows that this negative slope is highly significant (Table 1). On average, one additional microRNA target site leads to a 4% expression drop (i.e., a slope of ~ −0.06) in mammals. Since the observed slopes are somewhat smaller than the observed effect size from microRNA knock-out experiment, we simulate the slope by using the repression data obtained from the miRNA knockout lines to check the statistical significance. Such data are shown in Fig. 2 and the details are given in Methods. In most tissues, the observed slope is within the 95% confidence interval of simulations but is often on the low side. (Fig. 4A, 4C and 4D). In one tissue (Fig. 4B), the observed slope is significantly smaller than expected.

Importantly, the correlation is fairly low with R^2 < 0.05. The conclusion is the same when we weight every target site by the expression level of its governing miRNA. These results suggest that, in long-term expression evolution, miRNA target site changes contribute marginally to the gene expression divergence because the gains and losses often cancel out. It seems clear that small-step expression changes mediated by miRNAs do not lead to long-term directional changes. Such small-step evolution is nevertheless central to the continual fine-tuning near the optimum.

### 5. Modest contribution of small-step changes to the long-term phenotypic evolution

Fig. 4 shows that both the slope and the correlation between the expression divergence and target site changes are very low in long-term evolution. In this section, we will show that the low correlations, as well as the smaller-than-expected slopes, can be adequately explained. Even in such a simple model, many small-step changes can be overshadowed by a few large-step ones, as if the former do not matter in long-term phenotypic evolution.

Let us assume an initial expression optimum where the adaptive peak is. At times, the environment would change, resulting in a shift in the optimal expression level. Note that models of phenotypic evolution should, in principle, factor in environmental changes that drive peak shifts although empirical data on either the frequency or extent of such shifts are rare. If the new optimum after the environmental change is a distance from the starting point, large-step changes would be favored. For gene expression, large-step changes may be realized by transcription factors, promoters, enhancers and even chromatin structure. After the expression evolves to the vicinity of the new optimum, fine-tuning of gene expression by small steps is achieved by mutations in miRNA target sites. Fig. 5A depicts such a scenario which is simulated using the parameter values of Methods. The two panels show visually how large-step evolution followed by several small-step changes can be an efficient approach to a new adaptive peak.

**Figure 5.**
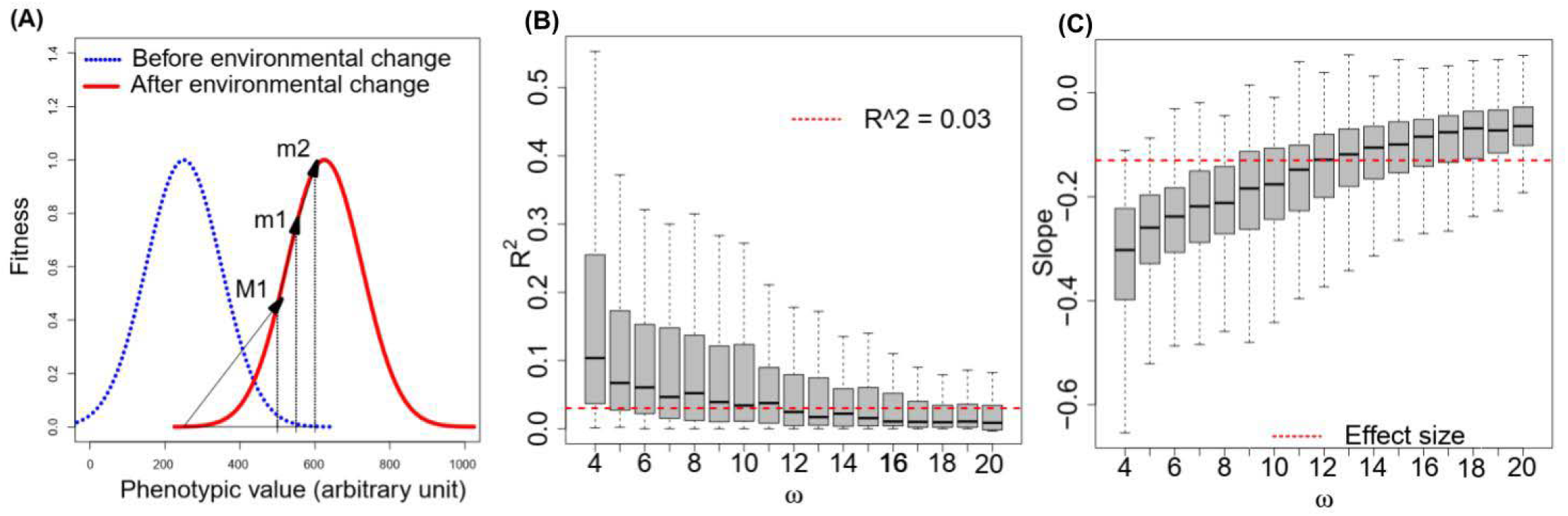
Simulation of gene expression evolution by separating large- and small-step changes after the same environmental shift in the optimal expression value. (A) one large-step change (M1) is followed by two small-step changes (m1 and m2). All changes are advantageous, and the optimum is gradually approached. (B) R^2^ by small step changes found in simulations. Observed R^2^ by miRNA target site (red dashed line) can be reproduced by a wide range of parameters. (C) Slopes by small step changes can deviate from the mechanical effect size (red dashed lines).

In simulating the approach to the new optimum, we focus on the width of fitness function, i.e., ω, which is negatively correlated with the efficiency of natural selection (see Methods). A small ω means that the phenotype has to be very close to the optimum to have high fitness, a very narrow fitness function. For example, let us consider a case where the distance of a phenotype to the optimum is 1 and an advantageous mutation shortens the distance by half. When ω = 5, the fitness advantage would be 0.015 but when ω = 15, it would decrease to 0.0016. At ω = 5, selection would be more efficient. We found that both the observed R^2 and slopes for miRNA target site changes could be well explained by a high level of ω.

Based on the simulations, Fig 5B shows that small-step changes could cumulatively explain 20% of the variation of expression evolution when ω was low, which is much higher than our observation on miRNA target site. As ω went higher, R^2 became lower and close to the observed level for miRNA target site (red dashed line in Fig. 5B). Fig 5C also shows that slopes of small-step changes are correlated with ω. In particular, slopes of small-step changes (evolutionary effect size) tend to be lower than the effect size of small-step changes (red dashed line in Fig. 5C) when ω was higher. These observations are consistent with our results on miRNA target sites.

In conclusion, Fig. 5 shows that large-step changes followed by many small-step adjustments can yield results compatible with the observed evolution in gene expression. Both the slope and the weak correlation of Fig. 4 can be obtained with reasonable parameter values in the simulations.

## Discussion

Changes in environmental conditions often demand phenotypic evolution and adaptive responses. For example, high-altitude adaptation in Tibet human population could be associated to specific mutations in genes related to hypoxic pathway (He et al. 2018). In another example, adaptation of woody plants after invading the intertidal zones consists of numerous expression changes (He et al. 2019; He et al. 2020). The question of whether phenotypic evolution proceeds by large or small steps must be examined in the context of environmental changes that would dictate the expression optimum. The FGM postulate may or may not be correct depending on several contextual attributes.

The first one is how often environmental changes result in large shifts in the phenotypic optimum. These changes should have been more likely in long term evolution between different genera or families than between congeneric species. If the environment has not changed in the recent past, one may reasonably assume that the population is near the optimum. Fisher proposed his solution of FGM in this context. For gene expression, the bulk of long-term divergence may have been carried out by large-step changes, but adaptive small-step changes are more numerous and likely fine-tuning in nature. It is worth noting that the environment does not only mean the physical one of temperature, nutrients, etc. In real life, Lenski and colleagues (Wiser et al. 2013; Good et al. 2017) have shown that natural selection continues in chemostats where the physical environment remains constant for two decades. It is possible that the genomic environment (i.e., gene interactions) never ceases to change.

The second attribute is the fitness landscape itself. In the landscape of FGM, all mutations are either beneficial or deleterious and small step changes are less likely to be deleterious. In the study of amino acid substitutions using the physiochemical distance as the phenotype, Chen and colleagues have found that large-step changes tend to be either more beneficial or more deleterious than small-step changes (Chen et al. 2019a, 2019b). However, if the mutations tend to lead to small-step phenotypic changes, then phenotypic evolution may still appear to be by small steps, even though large-step changes are more likely to be adaptive. For a resolution of the contrasting views on this issue, please consult Figure 3a in Chen et al. 2019a.

The third attribute concerns the genetic basis of the phenotypic trait. If the phenotype has a multi-genic basis, then it would require a series of mutations to reach the new optimum. The traditional view of gradual evolution may be along this line of thinking. Indeed, most phenotypes of interest are multi-genic in a very high dimension. These phenotypes may include sexual behavior (Hollocher et al. 1997a, 1997b), postmating fertility (Ting et al. 1998; Sun et al. 2004), and genital morphology (Hagen, et al. 2019). Compared to these phenotypes, gene expression level has a much simpler genetic basis when it is regulated by miRNAs. Note that a mutation in the miRNA target site vs. one in the binding site of the transcription factor would lead to different “step sizes” in the expression level.

The answer to the question of the small- vs. large-step phenotypic evolution has to be context-dependent. For example, the environmental factors, the shape of the fitness landscape and the genetic basis of the phenotype would all influence the answer. The answer also depends on whether the question is long-term or short-term evolution. For short-term evolution in the vicinity of the adaptive peak, FGM appears to be an appropriate model. Finally, while this current study is based on phenotypes with a well-defined genetic basis, a companion study (Lu et al. submitted) provides the functional reasons that phenotypic evolution will have to be modeled on the genetic basis of the phenotype being studied.

## Materials and methods

### Expression of coding genes in mammals and Drosophila

RNA-sequencing raw reads, originally produced by the Kaessmann lab, were downloaded from GEO (accession GSE30352) (Brawand et al. 2011). Read mapping was performed using TopHat (Kim et al. 2013) (version 2.1.0) with the Hg38 human and mm10 mouse gene annotation. Relative abundance for each transcript was estimated with Cufflinks (Trapnell et al. 2010) (version 2.1.1) using the FPKM method. If multiple isoforms of a gene were expressed in the same tissue, we retained the most abundant transcript. The R function normalize.quantiles, implemented in the preprocessCore (Bolstad 2018) package, was used to normalize the gene expression data. We have also applied a scaling procedure based on genes with most conserved ranks of expression between species, similar to the one used by Brawand et al (Brawand et al 2011), to normalize the expression values. Using this method does not alter our conclusions. Drosophila melanogaster and Drosophila yakuba mRNA sequencing raw reads were downloaded from GEO (GSE99574), gene expression levels for each coding gene are computed using the same method as described above. For Drosophila melanogaster, results based on gene expression of strain W1118 are presented. The conclusions remain the same when using the Oregon R strain. Genomic annotations of D. melanogaster and D.yakuba were downloaded from Flybase (version dmel_r6.26_FB2019_01 and dyak_r1.05_FB2018_06, respectively)

### microRNA expression in mammals and Drosophila

Small-RNA sequencing raw reads were originally produced by the Kaessmann lab and downloaded from GEO with accession GSE40499 (Meunier et al. 2013). Raw reads were processed using trim_galore (Krueger 2015) (version 0.4.1), removing the adapter sequence ATCTCGTATGCCGTCTTCTGCTTG. Processed reads were first aligned to human and mouse genomes (Hg38 and mm10) to determine the expression of each miRNA locus using Bowtie (Langmead et al. 2009) (version 1.1.2) with parameters “-v 0 -a --best --strata”, allowing no mismatches and reporting all alignments per read (restricting to hits guaranteed best stratum). We excluded reads that mapped to genomic regions outside of known microRNA loci. Next, processed reads were mapped to mature microRNA sequences (including 3’ and 5’ flanking 3 base-pairs) and the corresponding seeds counted using an in-house script. Thus, mature miRNA expression was collapsed into seeds (i.e., families). If a read aligned to multiple mature microRNA with different seeds, its counts were equally distributed. For Drosophila small RNA expression, preprocessed unique reads were downloaded from GEO (GSE12840, GSE11624 and GSE98013). Abundance for each miRNA locus and family are computed in the same way as for the mammalian data.

### Analysis of the microRNA knock-out transcriptome

We collected trancriptome data from mice and D. melanogaster with microRNAs knocked out. Raw data were downloaded from GEO, with accessions GSE41090, GSE60426, GSE45760, GSE97364, GSE118004 and GSE73946, corresponding to knockouts of mmu-miR-181 (Henao-Mejia et al. 2013), mmu-miR-122 (Eichhorn et al. 2014), mmu-miR-1 (Wei et al. 2014), dme-miR-310 (Zhao et al. 2018), dme-miR-210 (Weigelt et al. 2019) and dme-miR-11 (Truscott et al. 2016), respectively. We processed read mapping and gene expression quantification as described above. When multiple isoforms were present, the transcript with the highest expression in wild type was used. Target gene expression from replicates was averaged. Expression change of target genes and non-target background genes was compared using the two-sample Kolmogorov-Smirnov test. For mouse miRNA, target sites present in human and mouse are considered conserved. For D. melanogaster miRNAs, we considered target sites present in D. melanogaster and D. yakuba conserved, while those absent in D. yakuba as non-conserved.

### Estimating the contribution of miRNA target site change to gene expression evolution

TargetScan (Lewis et al. 2005; Agarwal et al. 2015) was used to scan a multiple species alignment of 3’UTR sequences (including human, chimp, macaque, mouse and rat) to identify target sites for the top 20 miRNA seeds. Only three canonical types of target sites were used (7mer-A1, 7mer-m8, and 8mer-A1). When using mouse miRNA knock-out data to infer repression strength of each target site, we used genes with only one target site regulated by the microRNA. A linear model was used to estimate the effect of a miRNA target site change on gene expression evolution. Observed target gene expression change was used as the response variable (log2 transformed) while the observed target site change was the predictor. Regression analysis was performed using the R function lm that implemented in the stats package.

### Expected gene expression change due to miRNA target site change

We predict gene expression divergence due to miRNA target site change in evolution for human and mouse. Briefly, observed target site changes (polarized Net(i)) for each gene was used and the corresponding target gene expression divergence was simulated by sampling target expression change from the KO experiment. The simulated gene expression divergence was then regressed on observed Net(i) to compute the simulated slope. For example, if a gene has Hi - Mi = 3, we sample three target genes from knock-out data. If these three random target genes are up-regulated 50%, 20% and 10% in KO relative to wild-type, we can infer that their expressions are repressed by 50%, 20% and 10% in vivo. We then predict that expression level of this gene with Hi - Mi = 3 may decrease 64% in human relative to mouse, i.e., 1 - (1 - 50%) × (1 - 20%) × (1 - 10%). Thus, expression change for each gene can be inferred, providing an expectation of expression change driven by microRNA target site change exclusively. This inferred expression change (log2 transformed) was then regressed on observed Hi - Mi using the R function lm. We ran target site sampling 1,000 times for each tissue to estimate 95% confidence intervals for both R^2^ and the slope. If the observed slope is higher than the simulated slope (observed target site effect in evolution is stronger than expectation), that suggests other regulatory changes during evolution predominantly work in the same direction as Net(i); if the observed slope is lower than simulated slope (observed target site effect in evolution is weaker than expectation), that suggests other regulatory changes during evolution predominantly work in the opposite direction from Net(i).

### The structure of the gene expression evolution model and simulations

To explain the observations that many small changes in gene expression do not lead to large divergence, we implemented a simulation where expression divergence proceeds by both large and small step changes. Large-step changes reflect the evolution of transcriptional regulatory elements (e.g., transcription factors, promoters, enhancers). In contrast, fine-tuning of gene expression by small steps is achieved by mutations in miRNA target sites. Our model assumes an initial state of expression stasis at the optimal expression level. After the environment changes (e.g. drought), a shift in the optimal expression level for gene *i* occurs. If a mutation drives the expression of gene *i* towards the new optimum, that mutation would be favored by natural selection.

Fig. 5A illustrates this model. In the left panel, mutation M1, a large-step change, occurred first to increase the phenotypic value towards the optimum. Several small step changes then provide further fine-tuning of the phenotype. In practice, we assume that shift of gene expression optimum caused by environmental change follows an exponential distribution:

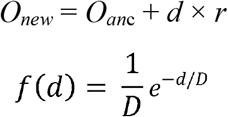

Here *O_anc_* and *O_new_* are phenotypic optima before and after the environmental change while *d* is the phenotypic distance between *O_new_* and *O_anc_*, which follows an exponential distribution with mean *D*. We also assume that *O_new_* can be either higher or lower than *O_anc_* and therefore set *r* to be a Bernoulli random variable, which could be 1 or −1, each with a probability of 50%. During the evolution towards a novel optimum, we assumed that the effect size of mutation falls into two classes: one represents large-step change while the other represents small-step change. Note that we chose a simple way to simulate this process and the assumption could be wild. However, it would be adequate to address the adaptive process in a framework of FGM.

Here, a random mutation would cause a large-step change with probability P_L_ and a small-step change with probability 1 - P_L_. The mutation effect sizes also follow an exponential distribution in our simulation, with the mean effect size of *B_1_* and *B_2_* for small steps and large steps, respectively.

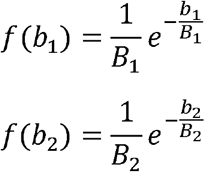

The sign of the effect of a mutation is also a Bernoulli random variable, which could be positive or negative, each with a probability of 50%. Once a mutation occurs and its effect size *b_k_* (k = 1,2) is determined, fitness of the mutant is compared to wild type. We model fitness using a Gaussian fitness function, which compares expression level of mutant and wild type with the novel optimum *O_new_* using equation (1) (Lande 1976).

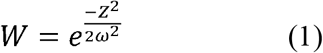

Here *ω* is the “width” of the fitness function. When *ω* is small, stronger purifying selection (Wang et al. 2018; Chen et al. 2021; Wen et al. 2021) is imposed on deviations from *O_new_*, denoted as *z*. Based on equation (1), we have:

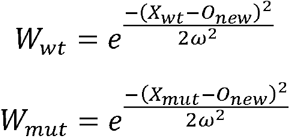

Here *X* is the expression level for mutant and wild-type. If *X_mut_* is closer to *O_new_* than *X_wt_*, the mutant confers higher fitness. For each mutant,

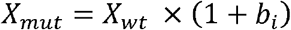

Where *i* could be 1 or 2, representing small-step or large-step change. When the environment has just changed and mutations have yet to occur, *X_wt_* is set to 0. Based on equation (1), fitness of both the wild-type and the mutant can be determined. The selective coefficient *s* for a mutation is then:

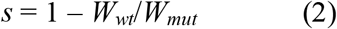

Finally, the fixation probability for each mutant (Kimura 1962; Crow and Kimura 1970) can be calculated as

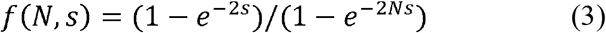

Once a mutation is fixed, *X_wt_* will be replaced by *X_mut_*, and *W_wt_* will be replaced by *W_mut_*. Otherwise, both *X_wt_* and *W_wt_* remain unchanged (Ruan et al. 2021; Ruan et al. 2022) The simulation would be terminated when the distance between a phenotypic state and the optimum was smaller than 50% of the mean of distribution of small-step changes. We choose this criterion as it will possibly prevent small-step changes from overshooting the optimum.

We simulated this process for 10,000 genes using the following parameters: population size N = 10,000; *P_L_*: from 0.4 to 0.6; *ω*: from 4 to 20; B_1_: 10%; B_2_: 200%, 400%, 800%; D: 200%, 400%, 800%. For each set of parameters, we repeat the simulation for 1,000 replicates. Computer codes for the simulation are available at https://github.com/linpei26/FGMicroRNA.

## Acknowledgements

We would like to thank Tian Tang and Rui Zhang for advice on data processing and analysis, Anthony J. Greenberg for advice on manuscript writing, anonymous editors and reviewers for their critical and insightful reviews. We thank all members in the Wu lab for helpful comments and sharing of ideas. This work was supported by National Natural Science Foundation of China (31730046,32150006,91731000) Innovation Group Project of Southern Marine Science and Engineering Guangdong Laboratory (Zhuhai) (No. 311021006), National Key Research and Development Projects of the Ministry of Science and Technology of China (2021YFC2301300, National Key R&D Program of China (2021YFC0863400).

