## Supplementary Note 1 for "Gene expression as phenotype - Many small-step changes leading to little long-term phenotypic evolution"

**A literature survey on the evolution of phenotypes in large vs. small steps in relation to Fisher’s Geometric Model (FGM)**

The idea that organisms evolve in very high-dimensional space leads to the widespread belief that small-step mutations should be the basis of adaption. Adaptive evolution in incremental small steps was formally encapsulated by R. A. Fisher’s geometric model (FGM). The essence of FGM is phenotypic evolution in small steps. Nevertheless, it is a specific view of adaptation that is distinguished from many similar views of “evolution in small steps”. For example, the gradual evolution in phenotype is also the hallmark of Darwinism. In the Darwinian view, each large-step phenotypic change usually requires multiple steps of evolution (or multiple mutations in the modern lexicon). From a pre-molecular biology perspective, the small steps are genetically and developmentally constrained by what mutations can bring about. In contrast, FGM is about the preference of natural selection for small-step changes, given the options of both large-step and small-step changes.

Small-step change is also a key rule of neutral molecular evolution (Kimura 1983, Chap. 4). In the neutralists’ view, natural selection is predominantly of the negative kind. Hence, the less deviant from the wild-type, the less deleterious the mutation would be. Again, this view is in contrast to FGM which is concerned mainly with positive selection. In this section, we survey the literature of the last two decades on the subject of phenotypic evolution in small vs. large steps, most but not all of which explicitly refers to FGM. The literature is roughly in three categories.

**Category I**. FGM has been widely utilized and extended by theoretical studies on adaptation (Orr 1998, 1999; Hietpas, et al. 2013; Lourenço, et al. 2013; Matuszewski, et al. 2014; Moura de Sousa, et al. 2016), speciation (Fraïsse, et al. 2016; Yamaguchi and Otto 2020), epistasis (Schoustra, et al. 2016; Hwang, et al. 2017), balancing selection (Sellis, et al. 2011; Connallon and Clark 2014a, b), deleterious mutation load (Martin and Lenormand 2006; McCandlish, et al. 2014; Stearns and Fenster 2016; Huber, et al. 2017) and population rescue upon the risk of extinction (Anciaux, et al. 2018; Anciaux, et al. 2019; Osmond, et al. 2020). Theoretical studies generally accept the validity of FGM, without evaluating whether FGM fits empirical observations.

**Category II**. Experimental evolution studies have the potential to test FGM with empirical data (Burch and Chao 1999). Surprisingly, mutation size is widely measured by its effect on fitness rather than phenotypes (Burch and Chao 1999; Perfeito, et al. 2007; Schoustra, et al. 2009; Sousa, et al. 2012; Trindade, et al. 2012; Berger and Postma 2014; Blanquart and Bataillon 2016). These studies are in fact about the nature of fitness mutations. They ask whether mutations tend to generate large or small fitness gains. In contrast, FGM is about the working of natural selection – Does natural selection favor small or large phenotypic changes?

Clearly, studies of either category are very different from this present study, which is to test the underlying assumptions of FGM directly. In this present study, we propose that gene expression can be considered as a phenotype for a test of FGM.

**Category III**. There is indeed an extensive literature on phenotypic evolution step size (Simpson 1953; Lande 1976, 1985; Gould 2007). Gaussian processes have been used to model the evolution of body mass (Freckleton, et al. 2003), gene expression level (Brawand, et al. 2011) and many other traits (Harmon, et al. 2010; Uyeda, et al. 2011; Arnold 2014; Barua and Mikheyev 2020). The statistical analyses can infer evolutionary jumps for phenotypes, such as body mass and endocranial volume in primates (Landis, et al. 2013), body size in anoles (Duchen, et al. 2017), morphology of the digestive tract of parrots (Duchen, et al. 2017).

When phenotypic evolution is used to test FGM, the same limitation inherent in the Darwinian view of gradual evolution re-surfaces. Large-step phenotypic changes often require many small incremental steps due to mutational constraints. In that case, FGM does not apply. In FGM, natural selection needs the options of both large- and small-step changes to operate on. Hence, the genetic basis of the phenotypic change needs to be known conduct a test of FGM.

Many studies have indeed tried to delineate the genetic basis of phenotypic divergence between species (Jeong, et al. 2008; Chan, et al. 2010; Frankel, et al. 2011; Prescott, et al. 2015; Kvon, et al. 2016; Raissig, et al. 2016; Wallbank, et al. 2016; Li and Fay 2017; Roscito, et al. 2018; Paolino, et al. 2020). Among these studies, an important role of regulatory change and its effect on gene expression is often identified. Major gene expression change can arise from single mutational leaps, for example, a 17-bp snake-specific deletion in the ZRS enhancer (Kvon, et al. 2016), or from multiple steps, such as substitutions in the E6 enhancer in D.Sechellia (Frankel, et al. 2011).

In general, delineating the genetic basis of phenotypic change has been a most challenging task in modern genetics. Complex traits and non-monogenic diseases offer ample evidence of the challenges. In our study, microRNAs provide a most common genetic basis for small step phenotypic changes. While the repression by miRNAs may contribute only a fraction of the total gene expression change, the molecular mechanism is common across many genes (> 30% of gene being directly governed by miRNA repression) (Liufu, et al. 2017; Zhao, et al. 2017; Lu, et al. 2018; Zhao, et al. 2018). In conclusion, expression changes governed by miRNAs provide many phenotypes that have a well-defined molecular mechanism and are thus ideal for testing FGM.

**Appendix**:

A full list of publications citing Fisher’s geometric model (FGM)

1	Orr, H. A. THE POPULATION GENETICS OF ADAPTATION: THE DISTRIBUTION OF FACTORS FIXED DURING ADAPTIVE EVOLUTION. Evolution 52, 935-949, doi:10.1111/j.1558-5646.1998.tb01823.x (1998).

2	Burch, C. L. & Chao, L. Evolution by small steps and rugged landscapes in the RNA virus phi6. Genetics 151, 921-927 (1999).

3	Orr, H. A. The evolutionary genetics of adaptation: a simulation study. Genetical research 74, 207-214, doi:10.1017/s0016672399004164 (1999).

4	Poon, A. & Otto, S. P. Compensating for our load of mutations: freezing the meltdown of small populations. Evolution 54, 1467-1479, doi:10.1111/j.0014-3820.2000.tb00693.x (2000).

5	Orr, H. A. The population genetics of adaptation: the adaptation of DNA sequences. Evolution 56, 1317-1330, doi:10.1111/j.0014-3820.2002.tb01446.x (2002).

6	Welch, J. J. & Waxman, D. Modularity and the cost of complexity. Evolution 57, 1723-1734, doi:10.1111/j.0014-3820.2003.tb00581.x (2003).

7	Orr, H. A. Theories of adaptation: what they do and don't say. Genetica 123, 3-13, doi:10.1007/s10709-004-2702-3 (2005).

8	Waxman, D. & Welch, J. J. Fisher's microscope and Haldane's ellipse. The American naturalist 166, 447-457, doi:10.1086/444404 (2005).

9	Haygood, R. Proceedings of the SMBE Tri-National Young Investigators' Workshop 2005. Mutation rate and the cost of complexity. Mol Biol Evol 23, 957-963, doi:10.1093/molbev/msj104 (2006).

10	Hine, E. & Blows, M. W. Determining the effective dimensionality of the genetic variance-covariance matrix. Genetics 173, 1135-1144, doi:10.1534/genetics.105.054627 (2006).

11	Martin, G. & Lenormand, T. A general multivariate extension of Fisher's geometrical model and the distribution of mutation fitness effects across species. Evolution 60, 893-907 (2006).

12	Orr, H. A. The distribution of fitness effects among beneficial mutations in Fisher's geometric model of adaptation. J Theor Biol 238, 279-285, doi:10.1016/j.jtbi.2005.05.001 (2006).

13	Waxman, D. Fisher's geometrical model of evolutionary adaptation--beyond spherical geometry. J Theor Biol 241, 887-895, doi:10.1016/j.jtbi.2006.01.024 (2006).

14	Waxman, D. Mean curvature versus normality: a comparison of two approximations of Fisher's geometrical model. Theoretical population biology 71, 30-36, doi:10.1016/j.tpb.2006.08.004 (2007).

15	Moorad, J. A. & Promislow, D. E. A theory of age-dependent mutation and senescence. Genetics 179, 2061-2073, doi:10.1534/genetics.108.088526 (2008).

16	Sella, G. An exact steady state solution of Fisher's geometric model and other models. Theoretical population biology 75, 30-34, doi:10.1016/j.tpb.2008.10.001 (2009).

17	Chevin, L. M., Martin, G. & Lenormand, T. Fisher's model and the genomics of adaptation: restricted pleiotropy, heterogenous mutation, and parallel evolution. Evolution 64, 3213-3231, doi:10.1111/j.1558-5646.2010.01058.x (2010).

18	Le Nagard, H., Chao, L. & Tenaillon, O. The emergence of complexity and restricted pleiotropy in adapting networks. BMC evolutionary biology 11, 326, doi:10.1186/1471-2148-11-326 (2011).

19	Lourenço, J., Galtier, N. & Glémin, S. Complexity, pleiotropy, and the fitness effect of mutations. Evolution 65, 1559-1571, doi:10.1111/j.1558-5646.2011.01237.x (2011).

20	Sellis, D., Callahan, B. J., Petrov, D. A. & Messer, P. W. Heterozygote advantage as a natural consequence of adaptation in diploids. Proc Natl Acad Sci U S A 108, 20666-20671, doi:10.1073/pnas.1114573108 (2011).

21	Chao, J., Ward, E. S. & Ober, R. J. Fisher information matrix for branching processes with application to electron-multiplying charge-coupled devices. Multidimensional systems and signal processing 23, 349-379, doi:10.1007/s11045-011-0150-7 (2012).

22	Razeto-Barry, P., Díaz, J. & Vásquez, R. A. The nearly neutral and selection theories of molecular evolution under the fisher geometrical framework: substitution rate, population size, and complexity. Genetics 191, 523-534, doi:10.1534/genetics.112.138628 (2012).

23	Sousa, A., Magalhães, S. & Gordo, I. Cost of antibiotic resistance and the geometry of adaptation. Mol Biol Evol 29, 1417-1428, doi:10.1093/molbev/msr302 (2012).

24	Trindade, S., Sousa, A. & Gordo, I. Antibiotic resistance and stress in the light of Fisher's model. Evolution 66, 3815-3824, doi:10.1111/j.1558-5646.2012.01722.x (2012).

25	Zhang, X. S. Fisher's geometrical model of fitness landscape and variance in fitness within a changing environment. Evolution 66, 2350-2368, doi:10.1111/j.1558-5646.2012.01610.x (2012).

26	Chabris, C. F. et al. Why it is hard to find genes associated with social science traits: theoretical and empirical considerations. American journal of public health 103 Suppl 1, S152-166, doi:10.2105/ajph.2013.301327 (2013).

27	Gordo, I. & Campos, P. R. Evolution of clonal populations approaching a fitness peak. Biology letters 9, 20120239, doi:10.1098/rsbl.2012.0239 (2013).

28	Hietpas, R. T., Bank, C., Jensen, J. D. & Bolon, D. N. A. Shifting fitness landscapes in response to altered environments. Evolution 67, 3512-3522, doi:10.1111/evo.12207 (2013).

29	Lourenço, J. M., Glémin, S. & Galtier, N. The rate of molecular adaptation in a changing environment. Mol Biol Evol 30, 1292-1301, doi:10.1093/molbev/mst026 (2013).

30	Velenich, A. & Gore, J. The strength of genetic interactions scales weakly with mutational effects. Genome Biol 14, R76, doi:10.1186/gb-2013-14-7-r76 (2013).

31	Weinreich, D. M. & Knies, J. L. Fisher's geometric model of adaptation meets the functional synthesis: data on pairwise epistasis for fitness yields insights into the shape and size of phenotype space. Evolution 67, 2957-2972, doi:10.1111/evo.12156 (2013).

32	Bank, C., Hietpas, R. T., Wong, A., Bolon, D. N. & Jensen, J. D. A bayesian MCMC approach to assess the complete distribution of fitness effects of new mutations: uncovering the potential for adaptive walks in challenging environments. Genetics 196, 841-852, doi:10.1534/genetics.113.156190 (2014).

33	Berger, D. & Postma, E. Biased estimates of diminishing-returns epistasis? Empirical evidence revisited. Genetics 198, 1417-1420, doi:10.1534/genetics.114.169870 (2014).

34	Blanquart, F., Achaz, G., Bataillon, T. & Tenaillon, O. Properties of selected mutations and genotypic landscapes under Fisher's geometric model. Evolution 68, 3537-3554, doi:10.1111/evo.12545 (2014).

35	Chevin, L. M., Decorzent, G. & Lenormand, T. Niche dimensionality and the genetics of ecological speciation. Evolution 68, 1244-1256, doi:10.1111/evo.12346 (2014).

36	Connallon, T. & Clark, A. G. Balancing selection in species with separate sexes: insights from Fisher's geometric model. Genetics 197, 991-1006, doi:10.1534/genetics.114.165605 (2014).

37	Connallon, T. & Clark, A. G. Evolutionary inevitability of sexual antagonism. Proceedings. Biological sciences 281, 20132123, doi:10.1098/rspb.2013.2123 (2014).

38	Foll, M. et al. Influenza virus drug resistance: a time-sampled population genetics perspective. PLoS Genet 10, e1004185, doi:10.1371/journal.pgen.1004185 (2014).

39	Martin, G. Fisher's geometrical model emerges as a property of complex integrated phenotypic networks. Genetics 197, 237-255, doi:10.1534/genetics.113.160325 (2014).

40	Matuszewski, S., Hermisson, J. & Kopp, M. Fisher's geometric model with a moving optimum. Evolution 68, 2571-2588, doi:10.1111/evo.12465 (2014).

41	McCandlish, D. M., Epstein, C. L. & Plotkin, J. B. The inevitability of unconditionally deleterious substitutions during adaptation. Evolution 68, 1351-1364, doi:10.1111/evo.12350 (2014).

42	Perfeito, L., Sousa, A., Bataillon, T. & Gordo, I. Rates of fitness decline and rebound suggest pervasive epistasis. Evolution 68, 150-162, doi:10.1111/evo.12234 (2014).

43	Tenaillon, O. The Utility of Fisher's Geometric Model in Evolutionary Genetics. Annu Rev Ecol Evol Syst 45, 179-201, doi:10.1146/annurev-ecolsys-120213-091846 (2014).

44	Connallon, T. & Clark, A. G. The distribution of fitness effects in an uncertain world. Evolution 69, 1610-1618, doi:10.1111/evo.12673 (2015).

45	Kang, M. et al. eQTL epistasis: detecting epistatic effects and inferring hierarchical relationships of genes in biological pathways. Bioinformatics (Oxford, England) 31, 656-664, doi:10.1093/bioinformatics/btu727 (2015).

46	Martin, G. & Lenormand, T. The fitness effect of mutations across environments: Fisher's geometrical model with multiple optima. Evolution 69, 1433-1447, doi:10.1111/evo.12671 (2015).

47	Ram, Y. & Hadany, L. The probability of improvement in Fisher's geometric model: a probabilistic approach. Theoretical population biology 99, 1-6, doi:10.1016/j.tpb.2014.10.004 (2015).

48	Schick, A., Bailey, S. F. & Kassen, R. Evolution of Fitness Trade-Offs in Locally Adapted Populations of Pseudomonas fluorescens. The American naturalist 186 Suppl 1, S48-59, doi:10.1086/682932 (2015).

49	Blanquart, F. & Bataillon, T. Epistasis and the Structure of Fitness Landscapes: Are Experimental Fitness Landscapes Compatible with Fisher's Geometric Model? Genetics 203, 847-862, doi:10.1534/genetics.115.182691 (2016).

50	Fraïsse, C., Gunnarsson, P. A., Roze, D., Bierne, N. & Welch, J. J. The genetics of speciation: Insights from Fisher's geometric model. Evolution 70, 1450-1464, doi:10.1111/evo.12968 (2016).

51	Kronholm, I. & Collins, S. Epigenetic mutations can both help and hinder adaptive evolution. Mol Ecol 25, 1856-1868, doi:10.1111/mec.13296 (2016).

52	Martin, G. & Roques, L. The Nonstationary Dynamics of Fitness Distributions: Asexual Model with Epistasis and Standing Variation. Genetics 204, 1541-1558, doi:10.1534/genetics.116.187385 (2016).

53	Moura de Sousa, J. A., Alpedrinha, J., Campos, P. R. & Gordo, I. Competition and fixation of cohorts of adaptive mutations under Fisher geometrical model. PeerJ 4, e2256, doi:10.7717/peerj.2256 (2016).

54	Schoustra, S., Hwang, S., Krug, J. & de Visser, J. A. Diminishing-returns epistasis among random beneficial mutations in a multicellular fungus. Proceedings. Biological sciences 283, doi:10.1098/rspb.2016.1376 (2016).

55	Stearns, F. W. & Fenster, C. B. Fisher's geometric model predicts the effects of random mutations when tested in the wild. Evolution 70, 495-501, doi:10.1111/evo.12858 (2016).

56	Harmand, N., Gallet, R., Jabbour-Zahab, R., Martin, G. & Lenormand, T. Fisher's geometrical model and the mutational patterns of antibiotic resistance across dose gradients. Evolution 71, 23-37, doi:10.1111/evo.13111 (2017).

57	Huber, C. D., Kim, B. Y., Marsden, C. D. & Lohmueller, K. E. Determining the factors driving selective effects of new nonsynonymous mutations. Proc Natl Acad Sci U S A 114, 4465-4470, doi:10.1073/pnas.1619508114 (2017).

58	Hwang, S., Park, S. C. & Krug, J. Genotypic Complexity of Fisher's Geometric Model. Genetics 206, 1049-1079, doi:10.1534/genetics.116.199497 (2017).

59	Anciaux, Y., Chevin, L. M., Ronce, O. & Martin, G. Evolutionary Rescue over a Fitness Landscape. Genetics 209, 265-279, doi:10.1534/genetics.118.300908 (2018).

60	Russ, D. & Kishony, R. Additivity of inhibitory effects in multidrug combinations. Nature microbiology 3, 1339-1345, doi:10.1038/s41564-018-0252-1 (2018).

61	Simon, A., Bierne, N. & Welch, J. J. Coadapted genomes and selection on hybrids: Fisher's geometric model explains a variety of empirical patterns. Evolution letters 2, 472-498, doi:10.1002/evl3.66 (2018).

62	Sroka, C. J. & Nagaraja, H. N. Odds ratios from logistic, geometric, Poisson, and negative binomial regression models. BMC medical research methodology 18, 112, doi:10.1186/s12874-018-0568-9 (2018).

63	Anciaux, Y., Lambert, A., Ronce, O., Roques, L. & Martin, G. Population persistence under high mutation rate: From evolutionary rescue to lethal mutagenesis. Evolution 73, 1517-1532, doi:10.1111/evo.13771 (2019).

64	Chen, Q. et al. Molecular Evolution in Large Steps-Codon Substitutions under Positive Selection. Mol Biol Evol 36, 1862-1873, doi:10.1093/molbev/msz108 (2019).

65	Chen, Q., Lan, A., Shen, X. & Wu, C. I. Molecular Evolution in Small Steps under Prevailing Negative Selection: A Nearly Universal Rule of Codon Substitution. Genome Biol Evol 11, 2702-2712, doi:10.1093/gbe/evz192 (2019).

66	Fraïsse, C. & Welch, J. J. The distribution of epistasis on simple fitness landscapes. Biology letters 15, 20180881, doi:10.1098/rsbl.2018.0881 (2019).

67	Scott, T. J. & Queller, D. C. Long-term evolutionary conflict, Sisyphean arms races, and power in Fisher's geometric model. Ecology and evolution 9, 11243-11253, doi:10.1002/ece3.5625 (2019).

68	Lavigne, F., Martin, G., Anciaux, Y., Papaïx, J. & Roques, L. When sinks become sources: Adaptive colonization in asexuals. Evolution 74, 29-42, doi:10.1111/evo.13848 (2020).

69	Osmond, M. M., Otto, S. P. & Martin, G. Genetic Paths to Evolutionary Rescue and the Distribution of Fitness Effects Along Them. Genetics 214, 493-510, doi:10.1534/genetics.119.302890 (2020).

70	Schneemann, H., De Sanctis, B., Roze, D., Bierne, N. & Welch, J. J. The geometry and genetics of hybridization. Evolution, doi:10.1111/evo.14116 (2020).

71	Tikhonov, M., Kachru, S. & Fisher, D. S. A model for the interplay between plastic tradeoffs and evolution in changing environments. Proc Natl Acad Sci U S A 117, 8934-8940, doi:10.1073/pnas.1915537117 (2020).

72	Yamaguchi, R. & Otto, S. P. Insights from Fisher's geometric model on the likelihood of speciation under different histories of environmental change. Evolution, doi:10.1111/evo.14032 (2020).
