## Supplementary tables and figures for "Gene expression as phenotype - Many small-step changes leading to little long-term phenotypic evolution"

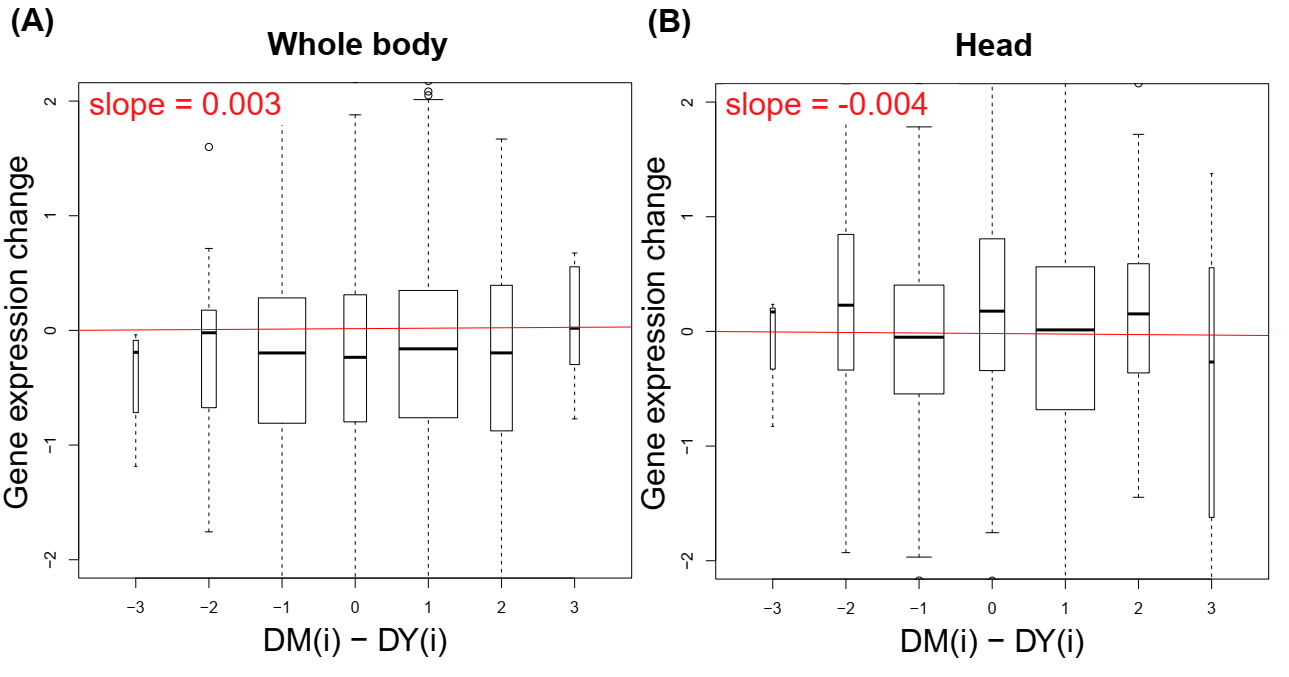

**Supplementary figure 1 . The accumulated effects of miRNA repression during drosophila evolution.** Distributions of expression changes of targets with different numbers of miRNA binding sites in six tissues (kidney, brain, heart, testes in primate and head, testis in Drosophila) during evolution. miRNA binding site changes of each gene are quantified by counting target sites present in one focal species but absent in the other closely-related species. Hi (Ri) stands for number of target sites on gene *i* that present in human (rhesus) but absent in rhesus (human), while DMi (DSi) stands for number of target sites on gene *i* that present in D.melanogaster (D.simulans) but absent in D.simulans (D.melanogaster). Red solid lines are drawn according to slopes computed by linear regression

|  | proportion of sequencing reads covered by top-20 miRNA families | number of miRNA families covering 95% of sequencing reads | proportion of sequencing reads covered by top-20 miRNA families | number of miRNAs families covering 95% of sequencing reads |
| --- | --- | --- | --- | --- |
|  | H.sapiens | | M. musculus | |
| brain | 92.45% | 30 | 85.33% | 48 |
| heart | 97.00% | 14 | 97.72% | 12 |
| kidney | 94.40% | 22 | 94.76% | 21 |
| testis | 91.90% | 29 | 94.18% | 23 |
|  | D.melanogaster | | D.yakuba | |
| Head | 95.1% | 27 | 89.5% | 33 |
| Whold-body | 90.2% | 30 | 87.7% | 32 |

Supplementary Table 1. Highly abundant miRNA families account for a large proportion of sequencing reads from the miRNA pool.

| microRNA | Target genes | number of target genes | mean | median | std.dev. |
| --- | --- | --- | --- | --- | --- |
| mmu-miR-223 | Mouse specific | 599 | 4.40% | 2.60% | 21.90% |
|  | conserved | 99 | 6.00% | 7.20% | 20.40% |
| mmu-miR-122 | Mouse specific | 586 | 24.10% | 22.50% | 46.10% |
|  | conserved | 62 | 55.20% | 55.30% | 47.90% |
| mmu-miR-181 | Mouse specific | 1275 | 0.10% | 5.40% | 51.50% |
|  | conserved | 400 | 11.00% | 14.80% | 43.70% |
| dme-miR-11 | D.melanogaster specific | 156 | 11% | 6.9% | 45.1% |
|  | conserved | 48 | 11% | 8.8% | 36.9% |
| dme-miR-310 | D.melanogaster specific | 262 | 1% | 0.3% | 68% |
|  | conserved | 152 | 5.3% | 1.7% | 72% |
| dme-miR-210 | D.melanogaster specific | 184 | 1.6% | 0.8% | 36% |
|  | conserved | 116 | 12% | 2.9% | 64% |

Supplementary Table 2. Distribution of repression effect size on target gene expression by conservative miRNAs.

|  |  | Net (i) | | Sum (i) | | Net (i) = 0 | |
| --- | --- | --- | --- | --- | --- | --- | --- |
|  |  | mean | Std.dev. | mean | Std.dev. | mean of sum(i) | Std.dev. of sum(i) |
| Mammal | Heart | 2.595 | 3.164 | 4.748 | 5.114 | 1.373 | 2.491 |
|  | Brain | 2.776 | 3.388 | 5.028 | 5.387 | 1.481 | 2.639 |
|  | Kidney | 2.571 | 3.141 | 4.690 | 5.072 | 1.356 | 2.509 |
|  | Testis | 2.893 | 3.566 | 5.481 | 5.957 | 1.580 | 3.068 |
| *Drosophila* | Head | 1.09 | 0.633 | 1.45 | 0.99 | 2.32 | 1.07 |
|  | body | 1.12 | 0.77 | 1.32 | 0.87 | 1.97 | 0.95 |

Supplementary Table 3. Summary statistics of Net(i) and Sum(i), based on top-20 miRNA families and target genes that expressing simultaneously in each tissue.
